# Modeling multifunctionality of genes with secondary gene co-expression networks in human brain provides novel disease insights

**DOI:** 10.1101/2020.09.29.317305

**Authors:** Juan A. Sánchez, Ana L. Gil-Martinez, Alejandro Cisterna, Sonia García-Ruíz, Alicia Gómez, Regina Reynolds, Mike Nalls, John Hardy, Mina Ryten, Juan A. Botía

## Abstract

**Motivation:** Co-expression networks are a powerful gene expression analysis method to study how genes co-express together in clusters with functional coherence that usually resemble specific cell type behaviour for the genes involved. They can be applied to bulk-tissue gene expression profiling and assign function, and usually cell type specificity, to a high percentage of the gene pool used to construct the network. One of the limitations of this method is that each gene is predicted to play a role in a specific set of coherent functions in a single cell type (i.e. at most we get a single <gene, function, cell type> for each gene). We present here GMSCA (Gene Multifunctionality Secondary Co-expression Analysis), a software tool that exploits the co-expression paradigm to increase the number of functions and cell types ascribed to a gene in bulk-tissue co-expression networks.

**Results:** We applied GMSCA to 27 co-expression networks derived from bulk-tissue gene expression profiling of a variety of brain tissues. Neurons and glial cells (microglia, astrocytes and oligodendrocytes) were considered the main cell types. Applying this approach, we increase the overall number of predicted triplets <gene, function, cell type> by 46.73%. Moreover, GMSCA predicts that the *SNCA* gene, traditionally associated to work mainly in neurons, also plays a relevant function in oligodendrocytes.

**Availability:** The tool is available at GitHub,https://github.com/drlaguna/GMSCA as open source software.

**Implementation:** GSMCA is implemented in R.

## Introduction

Gene co-expression networks (GCN) are a combination of gene clusters and graph networks, based on the correlation of mRNA levels from gene expression profiling [1]–[4]. Genes appearing together in a cluster or as neighbors in the network are said to be co-expressed. GCN analysis has proven to be a powerful tool for determining genes associated with molecular mechanisms underlying biological processes of interest and for defining the function of a gene in a cell type using bulk-tissue transcriptomic data. This approach provides insights into gene function in specific cell types by clustering co-expressed genes and detecting modules (i.e. gene clusters) enriched for cell type markers. By applying the “guilt-by-association” (GBA) heuristic [5], [6] to GCNs, all genes in a cell-type enriched module are then predicted to relate to a single cell type within which, they share the same function.

However, there is ample evidence to suggest that a single gene may have different biological functions in different cellular contexts. For example, the tumor suppressor gene, *TP53*, which encodes tumor protein p53 is described to have different roles depending on its interaction partners and consequently is implicated not only in DNA damage and repair, but also in the initiation of apoptosis and senescence [7]. Yet current GCN analyses cannot capture such complexity even if it is reflected in transcriptomic data because commonly used forms of GCN analysis assign a single gene to a single module. This is a significant limitation, especially when considering tissues with high cellular heterogeneity such as human brain tissue, where cellular context is likely to be of key importance to the understanding of gene function. Thus, elucidating the different roles of the mentioned gene, while accounting for different contexts, such as its expression in different cell types, could reveal new insights into the comprehension of gene function [8]–[11].

While cell-specific transcriptomic analyses may provide a means of addressing that issue, there are significant challenges associated with the construction of co-expression-based approaches in this context. Although single-cell technology is evolving rapidly, the gene expression data generated remains sparse in nature, with low sensitivity on the reads per gene and cell types detected when using single-cell and single-nucleus RNA-sequencing approaches, respectively. Other issues include drop-outs or transcriptional bursting (i.e. the time depending variation of transcription activity) [12]–[17]. Furthermore, cell-specific human brain gene expression data sets generated from large numbers of individuals are rare due to the associated costs, and are largely limited to single-nucleus RNA-sequencing (snRNAseq) data as tissue is primarily sampled from post-mortem tissue. These snRNAseq data sets not only tend to be particularly sparse, but also have systematic biases, such as the under-representation of genes expressed within the neuropil (i.e. the synaptically enriched area of the central nervous system). To overcome some of these challenges, alternative techniques have been developed to study specific cell types in bulk tissue, for example by deconvoluting the specific contribution of each cell type to gene expression [18]–[23].

Thus, the development of tools to maximize the value of the large quantities of deeply sequenced and publicly available human brain bulk-tissue transcriptomic datasets remains important. In this study, we develop a new method to investigate gene multifunctionality in bulk-tissue transcriptomic datasets. Here, we define multifunctionality as the association of a gene to multiple biological functions within a tissue as a result of its different cellular contexts. Particularly, we are primarily interested in the cell type context of genes. And our aim is to uncover different cell types and functions for the same genes and in this way opening new ways to study diseases with a genetic basis. We do this through the tool Gene Multifunctionality in Secondary Co-expression Network Analysis (GMSCA), which uses WGCNA-like co-expression networks to investigate gene multifunctionality on gene expression profiling from bulk tissue (see Figure 1a and b). GMSCA is applicable when single-cell based data is not available yet but also in settings where there are paired bulk-tissue and single-cell samples. Importantly, GMSCA has been developed not to focus on producing estimates of each cell contribution to gene expression but on predicting gene function in each cell type the gene is expressed in. This process involves two steps. Firstly, GMSCA constructs a primary gene co-expression network (PGCN) from a gene expression matrix to produce a primary set of triplets <gene, cell type, function> from the annotated PGCN. Secondly, GMSCA creates secondary gene co-expression networks (SGCN) for each of the target cell types. Briefly, for each cell type, and focusing on modules enriched for that cell type only as they appear in the PGCNs, GMSCA removes the contribution of that cell type from the expression matrix. GMSCA now constructs, with the newly created gene expression matrix (which includes all the original gene pool), a SGCN and extracts new triplets <gene, cell type, function> coming from it (see Figure 1c).

**Figure 1.**
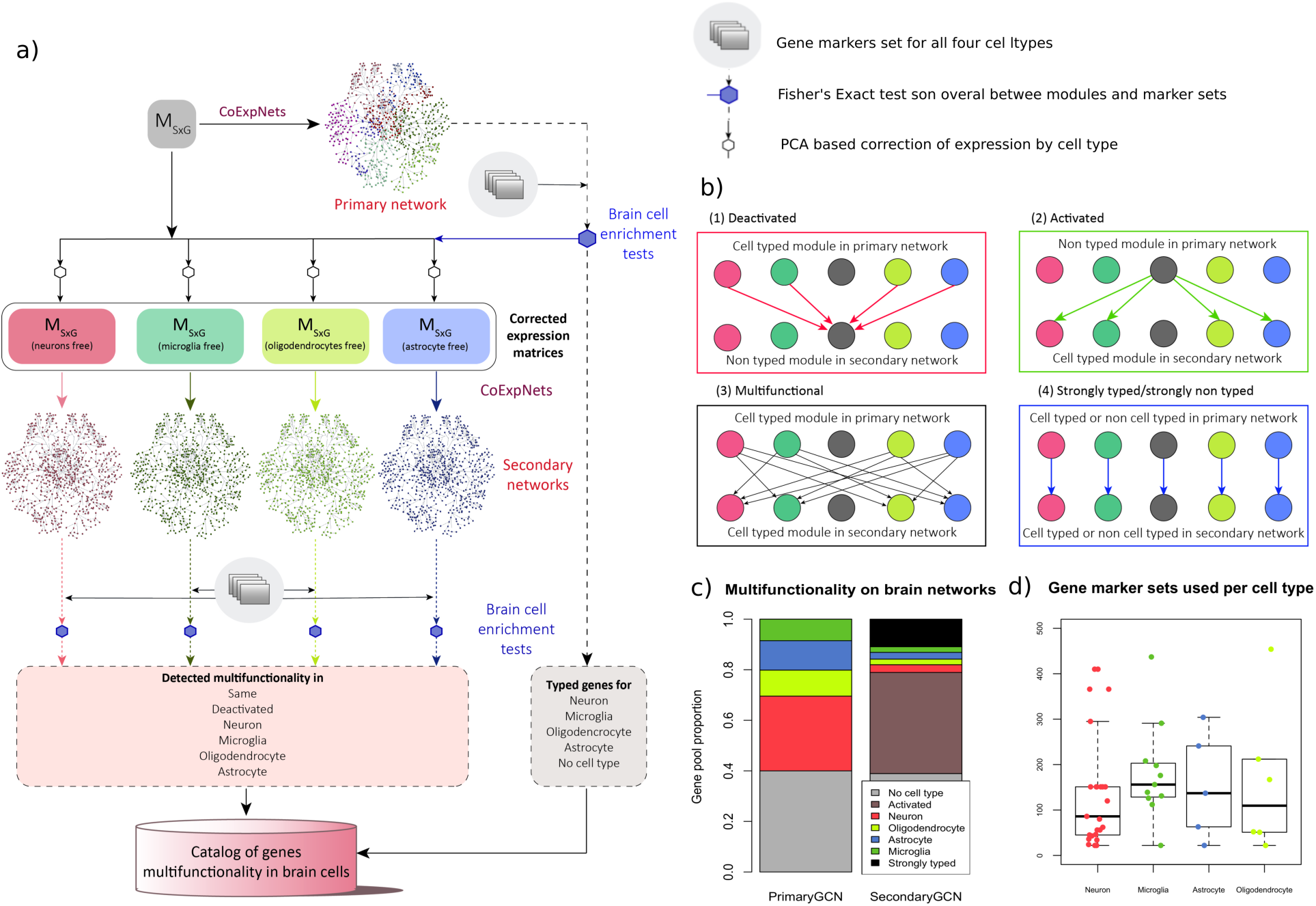
**a)** GMSCA generates a list of triplets <gene, cell type, function> for the genes included in the initial gene expression profiling matrix, M_SxG_. First, a WGCNA+k-means co-expression network called the primary network is created. Its modules are then tested (Fisher’s Exact Test, FET) for enrichment of cell type markers, in this case with brain cell type marker sets for neurons (in red), microglia (in green), oligodendrocytes (in light green) and astrocytes (in blue). Those modules with a clear signal (FET P < 0.05 on just a single cell type) are selected and their corresponding cell signal removed (see Methods) from expression to generate a new M’_SxG_. GMSCA creates a new co-expression network (these are called secondary) for each expression matrix and annotates their modules in the same way. Cell-type enriched modules in both primary and secondary networks generate as many triplets (gene, cell type, function) as genes in the module. **b)** Any gene found in a cell type enriched module is tagged by GMSCA as “typed”. When a gene in a primary co-expression network shifts from a cell-type enriched module to a non-cell-type enriched module in the secondary co-expression network, we say the gene is **deactivated** (red arrows). When it goes from a non-cell-type enriched module to a cell-type enriched module, we say the gene is **activated** (green arrows). A gene is **multifunctional** when it goes from a cell-type enriched module to a module with a different cell type enrichment (black arrows). A gene is **strongly typed** when it goes from a cell-type enriched module to another module with the same cell-type enrichment. It is **strongly non-typed** when the primary module and secondary module are both non-cell-type enriched. **c)** An average primary co-expression network tags 37% of genes as non-typed, another 30% as neuronal, 10% as oligodendrocytic, 13% as astrocytic and 8% as microglial. In a secondary network, 37% of non-typed genes become typed and 42.5% of typed genes become deactivated. Also, as 9.1% of the genes are tagged as pleiotropic in a single network, we can gain up to 36.4% annotations of pleiotropy with all four secondary networks GMSCA creates in this case. c) Number and size of gene markers set used by GMSCA for each cell type, in this paper.

GMSCA increases by 46.73% on average, the number of triplets obtained compared to conventional GCN, which notably increases its utility. Therefore, secondary co-expression networks can perfectly coexist with tissue deconvolution approaches as they generate additional functional annotation of different cell types. These new functional annotations include multiple annotations for single genes (i.e. they shape gene’s multifunctionality), creating new application avenues for this well-established analysis.

## Results

To investigate the utility of GMSCA for the prediction of gene multifunctionality in brain, we used 27 co-expression networks from three brain-derived gene expression datasets (see Table 1), including 13 bulk RNA-seq networks from GTEx’s [24] brain regions and 4 bulk RNA-seq networks from the ROSMAP [25], [26] project, all variations of frontal cortex samples. Further, we reused 10 bulk AffyExon Array networks derived from UKBEC’s 10 brain regions [27]. We refer to them as the GTEx, ROSMAP and UKBEC network families respectively.

**Table 1.**
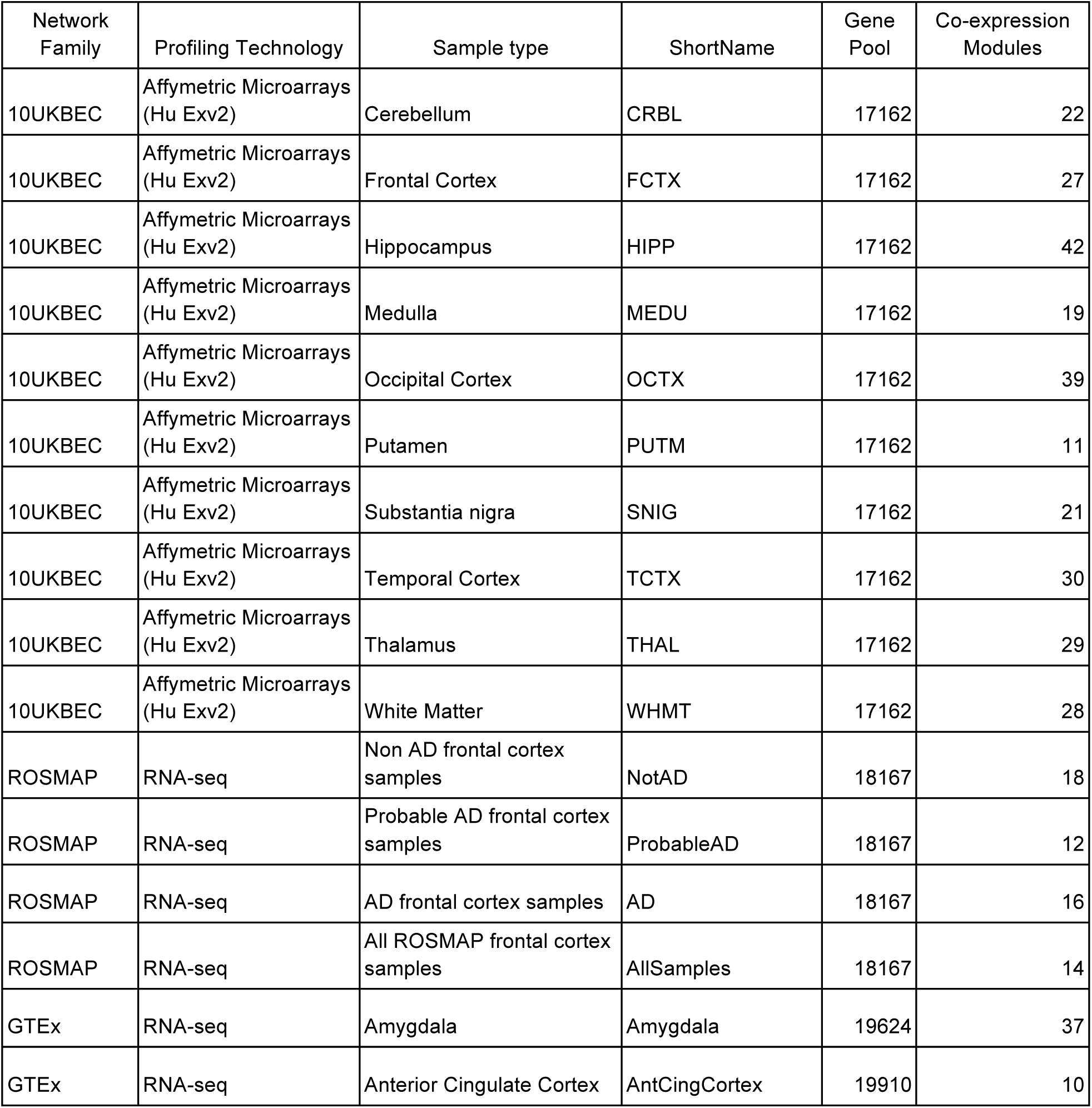
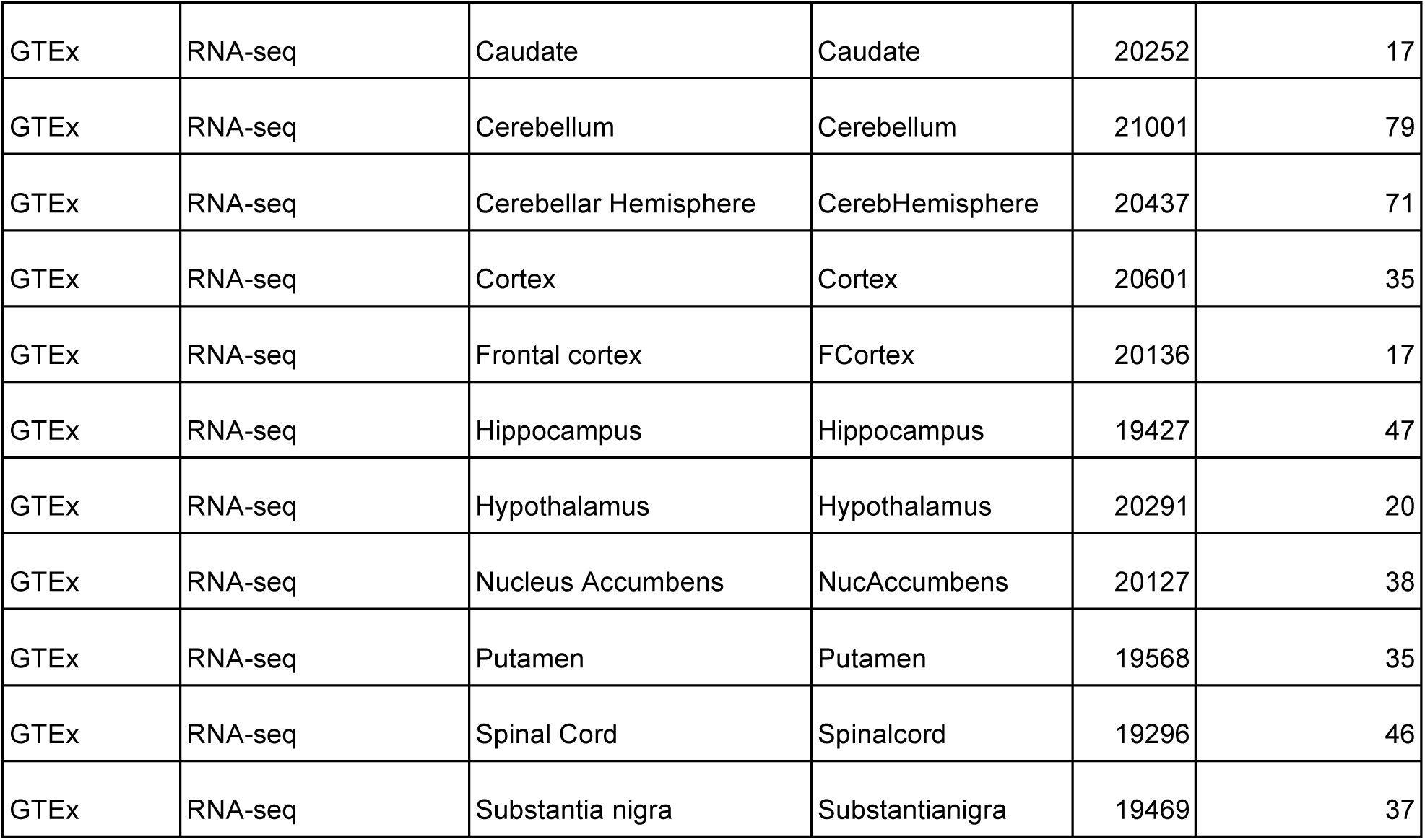
Networks used in the paper

The average gene pool for the three network families is 18682 and the average number of modules (clusters) is 30. As GMSCA uncovers multifunctionality, it focuses on modules enriched for markers of the cell types of interest. The average cell type enriched module size is 622 genes. Note that a percentage of those genes are cell type markers themselves; on average 158 genes, i.e. 25% of the whole module. Therefore, 75% of the genes in those modules generate new <gene, cell type, function> triplets in that cell type. Overall, the average PGCN generates more than 4500 of those prediction triplets. We focus on the major cell types in brain: neurons, microglia, astrocytes and oligodendrocytes (see Figure 1 d). We found that 44% of the PGCN modules enriched for markers of the major cell types were tagged as neurons, 21% as microglia, 18% as astrocytes and 17% as oligodendrocytes. Thanks to this approach, we gained the capability of studying gene multifunctionality by discovering new gene multifunctionality in specific cells with an additional 46.73% of the overall gene pool. 37% of the gene pool refers to newly activated genes and 9.73% are additional cell types for genes that become multifunctional (see Figure 1 c).

### GCN modules enriched for cell type markers are reliable

As all predictions made by GMSCA depend on cell marker enriched modules, we proceeded to assess whether those modules were replicable (i.e. preserved), and therefore reliable, across brain areas. The idea is that if a gene module is preserved in a similar tissue (i.e. post mortem brain tissue in this case), the module is credible and replicable. If the module is not preserved, it might be due to two reasons. Either the module is specific for that tissue, and hence reliable, or the module is not reliable, i.e. most genes are found there just by chance. We used WGNCA’s preservation analysis based on the estimation of a Z statistic called Z summary (see [24] for a detailed explanation). Values of Z over 10 for a module suggest strong module preservation. Values over 2 suggest some preservation. Modules with Z under 2 are not preserved at the tissue tested. See Figure 2 for a breakdown of test results across the three network families and cell types.

**Figure 2.**
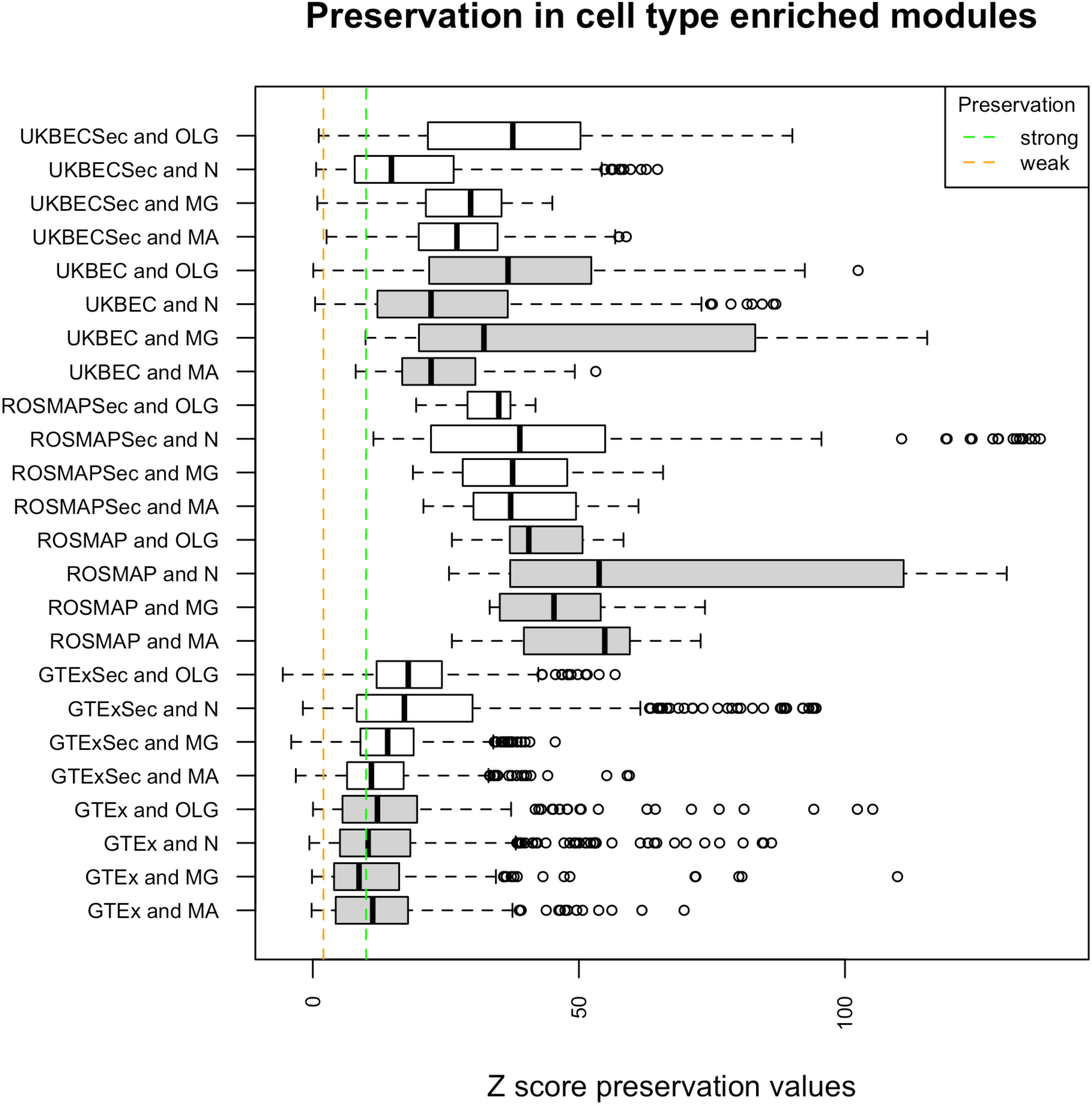
distribution of preservation values for all cell-type enriched modules as detected by GMSCA, for primary (in grey) and secondary (in white) networks. Each box plot corresponds to all modules within a family found to be enriched for the indicated cell type (MA for mature astrocytes, MG for microglia, N for neuron and OLG for oligodendrocytes). Families ending with “Sec” refer to secondary network modules. The vertical green dashed line marks the strong preservation limit (modules are easily replicable in other brain tissues and therefore highly reliable). The vertical orange dashed line marks the weak preservation limit (modules show signs or preservation and therefore some evidence of reliability). All preservation tests are performed within each network family. 98.1% of the preservervation tests sustain network reliability in UKBEC and 95.5% in GTEx. All tests yield strong preservation within the ROSMAP family.

We first tested GTEx PGCNs’s modules enriched for cell type-specific markers for preservation across the remaining GTEx PGCNs. This resulted in 560 preservation tests. We found that 8.6% of those tests were negative (i.e. Z < 2) suggesting that those modules were not preserved in the remaining brain PGCNs, while 40.4% of the tests showed signs of preservation (Z >= 2) and 50.8% showed strong preservation (Z >= 10). We performed the same analysis within the UKBEC networks. In this case, 630 tests were performed and 1.9% of the tests were negative. 13.3% pointed to some preservation and the remaining 84.7% indicated strong preservation. In the ROSMAP family, all target modules in all four networks are very well preserved. But this is something we may expect as one of the PGCNs includes all ROSMAP samples, and the other three include subsets of the samples used within that PGCN.

Next, we assessed the 52 GTEx SGCNs (i.e. each PGCN generates four new GCNs) for preservation. This resulted in 5157 tests being performed of which 4.4% were negative (a cell type enriched module was not preserved in a GTEx tissue), 25.9% implied weak preservation and 69.6% yielded strong preservation. We performed the same analysis for the 40 UKBEC SGCNs. In this case, 2310 preservation tests were performed and 1.12% of the tests were negative, whereas 17.7% of the tests showed signs of preservation and 81.12% showed strong preservation. All preservation tests in ROSMAP SGCNs showed strong preservation.

Part of the difference in non-preserved modules between GTEx and UKBEC can be explained due to PGCNs module size. The Z preservation value correlates positively with module size (R^2 0.38, supplementary Figure 1), i.e. bigger modules are more easily preserved [28]. Note that the maximum module size in any GTEx PGCN is 1663 whereas in UKBEC we find modules up to 2953 genes in size. Additionally, we wanted to further investigate whether the non-preserved GTEx modules could have had some biological meaning. For that purpose, we performed a gene level functional enrichment analysis on the genes at those modules based on the Gene Ontology, REACTOME and KEGG pathway databases using gProfileR R package [29]. On average, we obtained 164 annotation terms per non-preserved module, suggesting these modules were biologically relevant. Their lack of preservation may indicate that these modules were specific to their tissue.

### GMSCA multifunctional predictions replicate well in cell-type-specific datasets

GMSCA predicts gene multifunctionality in bulk-tissue RNAseq transcriptomics.. Therefore, replication of such predictions (at least when it comes to the cell type the gene is expressed in) in cell specific brain datasets would increase our confidence in the method. To evaluate this, we considered (a) two cell-specific gene expression datasets in human brain cortex, which are different in nature and (b) a mouse brain expression dataset [30] through the EWCE [31] tool.

We started with the replication of GMSCA predictions on human datasets. The first one is frontal cortex sc-ROSMAP [31]. It was obtained performing single cell transcriptomics in a fraction of bulk-RNAseq ROSMAP paired samples (24 AD cases and 24 controls). The second human dataset is based on immunopanning of temporal lobe cortex samples [32][33] from Barres Lab. This involved purification of specific cell types using cell surface markers followed by gene expression profiling on these purified cell types. As we already noted, sc-ROSMAP and Barres’ datasets are of different nature but both were reduced to a list of cell type specific genes for each of the cell types we are using in this paper (see methods). These lists are used then to compare with the cell multifunctionality predictions generated by GMSCA. Note that the genes used as cell markers within GMSCA are removed from the analysis in both bulk and single cell datasets to avoid optimistically biased results.

Multifunctionality predictions from all four ROSMAP PGCNs yielded significant overlap with the single nucleus ROSMAP dataset (see supplementary File 2). Focusing on the NotAD network (note that our sc-ROSMAP analysis includes non AD samples only), we obtained the following results for overlap Fisher’s Exact Test (FET): P < 4.27E-36 for neurons, P < 3.27E-46 for oligodendrocytes, 0.017 for microglia and 2.94E-42 for astrocytes. In regard to SGCN predictions, all overlaps are also significant, with FET P < 2.02E-36 for neurons, P < 2.4E-7 for oligodendrocytes, P < 2.3E-3 for Microglia and P < 1.28E-3 for astrocytes.

We assessed prediction replication in Barres Lab data by looking at how our cortex networks’ predictions replicated in Barres cell-specific gene expression profiles (see supplementary file 2) for all the results. As a way of example, we focus here on the same tissue used in Barres paper, temporal lobe (the 10UKBEC TCTX GCN). GMSCA generated highly significant overlaps in primary predictions with P < 4.51E-67 for neurons, P < 1.94E-31 for oligodendrocytes, P < 7.2E-13 for microglia and P < 1.01E-84 for astrocytes. Secondary predictions yielded significant overlaps for neurons (P < 1.19E-73), oligodendrocytes (P < 4.09E-42) and astrocytes (P 2.02E-12), but not for microglia (P 0.91).

Apart from human datasets, we also wanted to test replication of GMSCA predictions on different species through a more comprehensive single-cell brain transcriptomic dataset, with EWCE (Expression Weighted Cell Enrichment). This tool integrates mouse brain transcriptomics from 19 brain regions [47] and uses it as a reference. With EWCE it is possible to test whether a given gene list provided by the user presents higher expression levels in each cell type from the reference dataset than it would be expected by chance. We tested all cell type predictions from PGCNs and SGCNs to assess whether EWCE recapitulated similar enrichments and the analysis show that 100% of the GMSCA gene sets predicted to be functional in a given cell type are also found to be significantly enriched in expression by EWCE in the corresponding cell type (see methods, supplementary File 5)

### GMSCA multifunctional predictions show concordance across the three network families in cortex-like tissues

Thanks to the fact that we used different network families in this paper, we were able to compare same-tissue networks between families and assess the agreement level of predictions across their networks. Fortunately, we have cortex tissue networks across the three families. Specifically, we can find frontal cortex tissue in all three families, but also putamen and substantia nigra in GTEx and UKBEC (remind that ROSMAP is only frontal cortex). We assessed the significance of each pairwise overlap between (gene, cell-type) predictions from both primary and secondary networks, for each cell type and pair of frontal cortex tissues. All Fisher’s exact tests performed on the pairwise overlaps were highly significant, being the higher p-value of 1.3E-318 (see supplementary File 4 and supplementary figure 2).

In regard to the putamen brain area, 35% of the neuron predictions made from the UKBEC putamen networks (primary and secondary), are also found as neuron in the GTEx putamen networks. Note that the putamen UKBEC PGCN generates no microglia related predictions (it generates 2686 predictions in the SGCN though). The agreements for oligodendrocytes and astrocytes are 22.01% and 49.76%, respectively. For the substantia nigra brain area, the numbers for neuron, microglia, oligodendrocytes and astrocytes are 28.62%, 35.98%, 56.26%, and 42.91% respectively (see supplementary File 4 and supplementary figure 3).

### GMSCA links the *SNCA* gene to oligodendrocytes

The *SNCA* gene encodes the alpha-synuclein protein, located on the long arm of chromosome 4 at position 4q22.1. Mutations in this gene cause Parkinson disease (PD) [34]–[39]. *SNCA* is also linked to other neuro-degenerative diseases, mainly Alzheimer’s disease [40]. The alpha-synuclein protein is involved in the regulation of neurotransmitter release, synaptic function and plasticity of dopaminergic neurons[41]. Mutations in *SNCA* are associated with neuronal dysfunction. Therefore, it has been traditionally seen as highly relevant to neurons. Thanks to applying GMSCA to our networks, the generated multifunctionality predictions included an association of *SNCA* to oligodendrocytes in 4 SGCNs, see below (supplementary File 1).

*SNCA* is predominantly expressed in brain tissue (Figure 3 a), according to GTEx control samples bulk expression data V8 [24]. Moreover, *SNCA* is mainly expressed in neurons and oligodendrocytes [33] in the human cortex (Figure 3 b-c), as the Barres Lab data shows. In fact, *SNCA* was found in neuron-enriched modules in 10 out of the 13 GTEx PGCNs, all 4 ROSMAP networks and in 8 out of the 10 UKBEC networks. The only alternative cell type linked to *SNCA* is oligodendrocytes through the GTEx SGCNs of substantia nigra, putamen and hippocampus and the UKBEC temporal cortex (TCTX). All this data points to new avenues of research for this very important gene in neuro-degeneration.

**Figure 3.**
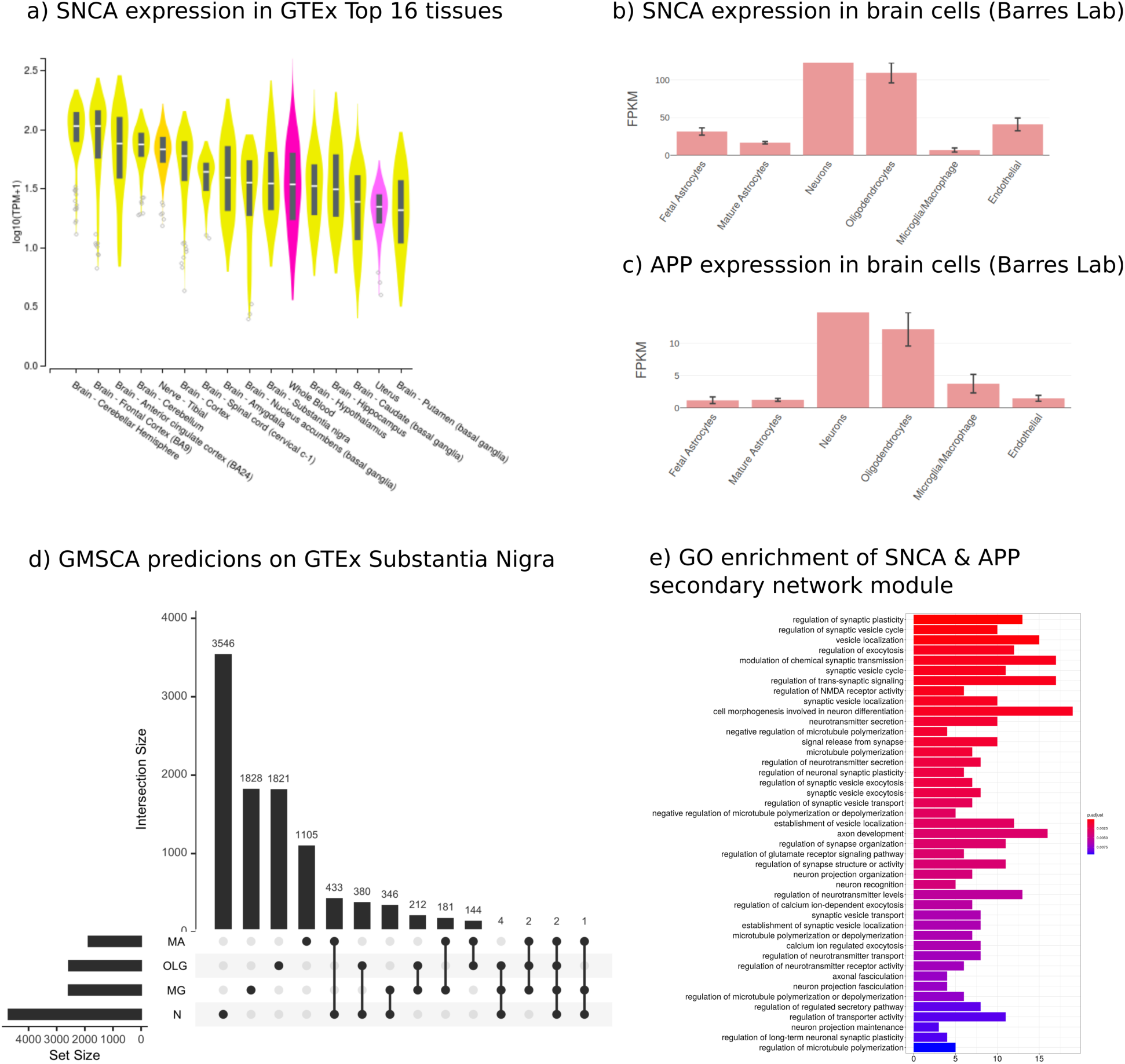
a) SNCA is predominantly expressed in brain, as the violin plots show. All 13 brain tissues are within the top 16 GTEx tissues expressing SNCA. b) Immunopanning data on brain cell expression from Barres Lab shows SNCA predominantly expresses in neurons and oligodendrocytes. GMSCA tags SNCA as neuronal and oligodendrocytic in the substantia nigra, putamen and hippocampus. c) APP, genetically linked to Alzheimer, is another interesting gene found in the same module as SNCA. Barres Lab’s data confirms it is predominantly expressed in neurons and oligodendrocytes. GMSCA tags it as neuronal and oligodendrocytic. d) UpSet plot (UpSet R package) on the genes from the predictions by GMSCA on the GTEx substantia nigra samples. SNCA and APP are found at the intersection between N (neurons) and OLG (oligodendrocytes) with 378 genes more.

The substantia nigra is a central tissue in PD. Therefore we focused the subsequent analysis on this tissue (Figure 3 d). *SNCA* is found in the orangered2 module of the PGCN (module membership, MM = 0.84). That module is enriched for dopaminergic neuron markers (P < 4.59E-25) and GO terms like catecholamine and dopamine biosynthetic processes (GO:0042423 P < 1.1E-5; GO:0042416 P < 9.9E-5), locomotory behavior (GO:0007626 P < 4.1E-4), neurotransmitter transport (GO:0006836 P < 1.9E-5), axonogenesis (GO:0007409 P < 6.4E-4), dopaminergic neuron differentiation (GO:0071542 P < 8.0E-3) and ferric ion transport (GO:0015682 P < 4.6E-4). Dopaminergic neuronal loss is the hallmark of PD. Moreover, the aforementioned biological processes are highly relevant to PD (Sawada, Oeda and Yamamoto 2013; Aggarwal, Reichert and VijayRaghavan 2019; Nutt, Carter and Sexton 2004; Song and Lee 2013; Hirsch 2009; Visanji et al. 2013). Interestingly, the darkorange2 module includes six more genes linked to PD (supplementary table 1). All this evidence suggests this network module is linked to PD.

When we applied GMSCA to create the corresponding SGCN, *SNCA* is found in the substantia nigra midnightblue module (MM = 0.53), which is enriched for markers of oligodendrocytes (P < 1.48E-4). This finding is replicated using the putamen and hippocampus co-expression networks, belonging to the same network family, GTEx. Moreover, themidnightblue module shared 27 genes with the *SNCA* SGCN module in putamen (Fisher’s exact test on the overlap significant, P < 5.3E-11) and 45 with the *SNCA* SGCN module at the hippocampus, P < 2.2E-16, suggesting function similarity of these modules across the three tissues linking *SNCA* to oligodendrocytes. The midnightblue module (Figure 3 e) was enriched for the following GO terms: synaptic plasticity (GO:0048167 P < 1.3E-4), neuronal plasticity (GO:0048168 P < 1.9E-3), regulation of synapse organization (GO:0050807 P < 3.5E-3), axonal fasciculation (GO:0007413 P < 7.2E-3), neuron projection fasciculation (GO:0106030 P < 7.2E-3), axon development (GO:0061564 P < 3.5E-3), neuron recognition (GO:0008038 P < 4.1E-3), regulation of neurotransmitter levels (GO:0001505 P < 5.5E-3), microtubule polymerization (GO:0046785 P < 1.4E-3), regulation of glutamate receptor signaling pathway (GO:1900449 P < 3.6E-3) and synaptic vesicle exocytosis (GO:0016079 P < 2.1E-3). Note that BP terms like “regulation of neurotransmitter levels” and “synaptic vesicle exocytosis” are processes found in the orangered2 module of the substantia nigra PGCN. Terms specific to the SGCN module, and also linked to oligodendrocytic activity, were: regulation of synapse organization (Eroglu and Barres 2010), regulation of glutamate receptor signaling pathway (Gautier et al. 2015), axon development (Simons and Nave 2016), axonal fasciculation, neuron projection fasciculation, neuron recognition or microtubule polymerization (Lee and Hur 2020). Interestingly, the midnightblue module included *APP* (MM = 0.84). Mutations in *APP* increase the risk of Alzheimer’s disease (Mullan et al. 1992). *APP*’s expression pattern in specific brain cells is similar to *SNCA*’s pattern, mainly expressed in neurons and oligodendrocytes (see Figure 3 c). The *APP* gene has been reported to have a role in regulating axonal myelination in oligodendrocytes [42].

A gene based analysis of the phenotypes linked to the midnightblue module (see methods) uncovered HPO terms like Dementia (HP:0000726 fold change 2.55), Myoclonus (HP:0001336 FC 2.13), Variable Expressivity (HP:0003828 FC 1.85), Anxiety (HP:0000739 FC 1.65), Intellectual severe disability (HP:0010864 FC 1.56), Depressivity (HP:0000716 FC 1.24), Tremor (HP:0001337 FC 1.22), Ataxia (HP:0001251 FC 1.2) and Dystonia (HP:0001332 FC: 1.16).

## Discussion

GCN-based analysis is a powerful tool for determining genes associated with molecular mechanisms, but its effectiveness is limited to assigning one gene to one function (i.e. a gene belongs to a specific module which, in turn, is annotated with a specific function). In a more realistic context, the biological environment is highly complex and a single gene may have different molecular functions, which it is described as multifunctionality [7]. This fact is especially relevant when studying cellular heterogeneity in human brain tissue, since one gene may have different functions depending on the cell type it is expressed in.

To overcome this issue, we propose GMSCA (Gene Multifunctionality Co-expression Analysis). GMSCA is a method to uncover additional co-expression profiles for genes, apart from those we obtain with conventional co-expression analyses. GMSCA significantly enhances the power of GCN analyses, delivering the capability to study gene multifunctionality in specific cells, as models of gene co-expression in bulk tissue. Fortunately, GMSCA needs the same inputs as conventional co-expression analysis on bulk tissue such as gene expression profiles, gene marker sets for the cell type under study and gene sets enrichment analysis tools as gProfileR for pathway or GO based module annotation. In this work, we used data from brain samples. Therefore, the SGCN obtained were focused on neurons, microglia, astrocytes and oligodendrocytes cell types. Interestingly, this approach increased the number of predictions of gene multifunctionality in specific cells with an additional 46.73% of the overall gene pool.

To demonstrate the potential of this method, we firstly assessed the reliability of GMSCA prediction triplets by means of its application to three network families for three different transcriptomics-based projects, UKBEC, GTEx and ROSMAP using WGCNA’s preservation analysis. We demonstrated that the cell-type enriched modules that GMSCA used to generate multifunctionality predictions were stable, including those in SGCNs. Part of the difference we obtained in non-preserved modules between GTEx and UKBEC could be explained by PGCNs module size.

Secondly, we showed a high level of replication of predictions in cell-type specific external data sets. As we already noted, sc-ROSMAP and Barres’ datasets are of different nature but both were reduced to a list of cell type specific genes for each of the cell types. Multifunctionality predictions from all four ROSMAP PGCNs yielded significant overlaps with the single nucleus ROSMAP dataset. We assessed prediction replication in Barres Lab data by looking at how our cortex networks’ predictions replicated in Barres cell specific gene expression profiles. All cell types but microglia showed highly significant levels of agreement. Moreover, thanks to the EWCE tool we were capable of assessing GMSCA predictions on mouse single-cell transcriptomics. 100% of both PGCN and SGCN gene sets yielded significant expression enrichment in equivalent brain cells.

Then, we tested the level of agreement in GMSCA predictions on the same tissues across the three networks families (frontal cortex tissue across the three families and substantia nigra and putamen tissues across GTEx and UKBEC) and demonstrated that agreement was high. We compared same-tissue networks between families and assessed the agreement level of predictions for cell marker enriched modules across their networks, in PGCNs and SGCNs, for preservation across brain areas.

We used this SGCN to get details about the biological function of *SNCA* within specific cell types, such as how relevant *SNCA*was in the corresponding module and what genes *SNCA* co-expressed with, e.g. APP linked to dementia and Alzheimer. In this study, *SNCA* was found in oligodendrocytes through the GTEx SGCNs of substantia nigra, putamen and hippocampus and through the UKBEC SGCNs of temporal cortex (TCTX). The relationship of *SNCA* and oligodendrocytes is supported by previous studies that have observed alpha-synuclein-containing inclusions (“coiled bodies”) in oligodendrocytes in parkinsonian brains [43]. Recently, the link between oligodendrocytes and Parkinson’s disease was reinforced integrating GWAS results with single-cell transcriptomic data [10] through testing the genes tagged by the GWAS outcome into the EWCE tool. Thus, GMSCA holds great potential to shed light on the cell types that genes are linked with and, in turn, how these genes may relate to molecular processes in disease.

In summary, GMSCA is a valuable strategy for characterizing the molecular functions of genes in specific cell types. This information is crucial to understanding the underlying mechanisms related to the initiation and progression of complex neurodegenerative disorders, such as Alzheimer’s disease and Parkinson’s disease. We are aware that some limitations still affect the sensitivity of this methodology, namely the brain sample acquisition or the limited availability of multi-omics data from the same individuals, but this method will help to improve the knowledge related to disease condition.

GMSCA is open source software, available at GitHub: https://github.com/drlaguna/GMSCA.

## Methods

### Co-expression network generation

For the purpose of this paper, we created three network families in the form of R packages as resources. These packages are CoExp10UKBEC, CoExpROSMAP and CoExpGTEx. Networks were created with WGCNA and refined with k-means with CoExpNets R package [1]. All the PGCNs are published at the CoExp Web site [44].

CoExp10UKBEC is a suite of networks created from confirmed non pathological human brain samples of 10 different brain areas of interest for neuro-degenerative diseases. Samples were profiled for gene expression with Affymetrix Human Exon v2. Microarrays. Identical gene pool was used to construct co-expression networks and details on initial WGCNA network construction can be found here [45]. Basic networks were refined with an additional step based on the k-means algorithm [1] implemented in theCoExpNets package.

ROSMAP is a transcriptomics oriented human frontal cortex resource which includes RNA-seq based gene expression profiling for 640 samples. Samples were corrected for batch effect using ComBat, and then age, sex, RIN and PMI (Post Mortem Interval) were regressed out of individual gene expression. We used the residuals to construct co-expression networks. We organise the 640 samples in four different groups to construct four different networks: one with all samples and three other networks arranged depending on neuritic plaques deposition postmortem diagnostic (a group for values 1, a second group for values 2 and 3 and a third group for values 4) with 200, 158, and 221 respectively. Networks are created with the CoExpNets method with a first step based on WGCNA and a refinement step based on k-means. All ROSMAP networks used in this paper can be found at the CoExpROSMAP R package.

We followed the same process for GTEx V6 gene expression. We focused on the development of 13 co-expression networks for the 13 available brain areas in GTEx RNA-seq samples. All samples are from bulk tissue in human control brains. We corrected samples for batch effect by using ComBat. We generated surrogated variables to model unknown effects in data and regressed those, age, sex, PMI and RIN from the overall gene expression. We used those residuals to construct the networks. All networks were created with the CoExpNets R package. All these networks from GTEx samples can be found at the CoExpGTEx package.

Note that, in all the three packages, once the expression matrices are ready for co-expression network generation, the procedure for creating a network (let it be either primary or secondary) is always the same. We find the right smooth parameter to satisfy scale free topology, then we generate an adjacency matrix and the subsequent topology overlap matrix (TOM). The next step is constructing a gene dendrogram using, as distance amongst genes, the matrix 1-TOM. Gene clusters are obtained by the cuTree algorithm with 100 as the minimum number of genes. We then apply a refinement step based on the k-means algorithm with 50 iteration steps. Genes are ranked within the modules through the Pearson correlation of their expression and the 1st principal component of gene expression of the whole module. This value is what is called module membership, MM. Finally, the gene clusters are annotated with function using gProfiler (see details here [1]). All networks are annotated for cell type as follows.

### Module Annotation for function and cell type

Each co-expression network clusters its gene pool into groups of genes that co-express together. The CoExpNets package was used to annotate all networks with functional annotations based on manually curated annotation databases: the Gene Ontology, REACTOME and KEGG. The gProfileR package is used for that.

In order to annotate co-expression network gene groups (modules) for specific brain cell types, we test each module for brain specific gene markers for the four main brain cell types (neuron, microglia, astrocytes and oligodendrocytes, see Figure 1 d) using manually curated cell marker datasets found at the CoExpNets package. We redesigned the original Fisher’s Exact test employed in CoExpNets to assess the significance of the overlap between genes at modules and genes at marker sets. This redesign was motivated by the detection of a significant association between network module size and -log10() transformation of p-values from the tests in all PGCNs, mean R^2^ 0.28, min 0.17, max 0.45 (supplementary Figure 4 a, b) and a subsequent inflation of the obtained p-values. Therefore, as well as applying Bonferroni correction, to account for multiple testing, we regress out gene set size effect from p-values significance (supplementary Figure 4 c, d) reducing the number of positive tests from an average of 22 per network to just a mean of 11.03.

### Gene Ontology and Human Phenotype Ontology Enrichment Analysis

The identification of enriched Gene Ontology (GO) terms in each gene module set was performed with the Bioconductor R package clusterProfiler [29]. The enrichGO function was used with Benjamin-Hochberg as a *p*-value (cutoff 0.01) correction method for the multiple testing and a *q*-value cutoff of 0.05.

Human Phenotype Ontology (HPO) terms associated with each human gene were downloaded from HPO [46]. To determine the enriched Human Phenotype Ontology (HPO) terms we have selected the relevant overrepresented phenotypes in each module, these are phenotypes that are in at least 2% of the genes from the module and Fold Change (FC) is a measure of how many times that phenotype is more likely to be found in the module than by chance (HPO ontology)

### Secondary co-expression networks generation

After generating the PGCN of a gene expression matrix, GMSCA detects modules enriched for the cell types of interest. For each gene in a module enriched for cell type ct, GMSCA generates a prediction triplet <gene, ct, function> where *function* refers to the Gene Ontology based annotation for all genes in the module. GMSCA uses PGCNs as the starting point for the generation of prediction triplets. But its main aim is to (1) discover additional cell types and functions for the already typed genes and to (2) uncover new cell types and functions for genes in non-enriched modules. This is possible thanks to **secondary co-expression networks**. GMSCA applies a transformation to the gene expression of genes at modules enriched for the cell types of interest. We model gene expression for any gene g as follows 

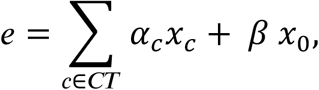

where e is the gene expression for a specific gene, CT is the set of all cell types considered of interest for the tissue, x_c_ is the particular contribution per sample of cell type c to the gene, and x_o_ represents other unknown factors contributing to the expression. GMSCA assumes this additive model to decompose the gene expression matrix of each cell type enriched module as follows. Let us suppose that module m is enriched for a particular cell type. GMSCA removes the contribution of that cell type to gene expression from module m as follows: it starts with the M_S,m_ matrix (i.e. gene expression of genes in module m for all samples S), it then obtains the set of PCAs to explain 90% of the variance, and compounds a new matrix with them, T_m,m,_, the transformed gene expression matrix. It then removes the 1st PCA from that matrix to create T’_m-1,m-1_. Note that the 1st PCA is assumed to be the cell type contribution to gene expression that we remove. T’_m-1,m-1_ is finally used to reconstruct the original expression, using all PCA axes but the 1st. The current matrix, M’_S,m_, is now free of the detected cell type contribution. This matrix is then used by GMSCA to create a new GCN that we call secondary, i.e. SGCN. GMSCA constructs, in this way, one SGCN for each cell type. To conclude this part, note that it is extremely rare to find gene modules enriched for more than one cell type. And in those cases, the enrichment of one of the cell types is almost marginal, while the other is highly significant. GMSCA drops out the marginal signal.

### Cell-type specific gene expression estimates for Barres

We downloaded supplementary table 4 from [33] which contains expression levels for neurons, mature astrocytes, microglia and oligodendrocytes. Within each cell type, we averaged all available values for each gene. We scale and log transform that matrix. Then we generate a boolean matrix of specific gene expression in cell types, M_g x ct_ with genes in rows and cell types in columns such that M[i,j] is set to TRUE when gene i has greater expression than the mean expression from the whole expression matrix.

In order to assess the multifunctionality prediction overlap between Barres data and our co-expression networks, we first look at GMSCA predictions coming from PGCNs, remove GMSCA marker genes from both GMSCA and Barres’, and assess the overlap in the corresponding cell type at the Barres’ data. We report a Fisher’s exact test on this overlap. Then we look at secondary predictions and report a Fisher’s exact test on the overlap. In each comparison, we remove known gene markers to avoid optimistic and unfair overlap assessments.

### Cell-type specific gene expression estimates for sc-ROSMAP

We downloaded the sc-ROSMAP filtered count matrix from the Synapse Web site. In sc-ROSMAP, we can find 48 samples of human frontal cortex tissue. Out of those, exactly 24 were given a diagnosis of Alzheimer’s disease cases and 24 controls. For this paper, we just kept the controls to avoid any potential bias due to neuro-degenerative or inflammatory processes. We dropped all zero count cell samples and generate a boolean matrix of gene expression in neurons, microglia, mature astrocytes and oligodendrocytes, M_g x ct_ with genes in rows and cell types in columns such that M[i,j] is TRUE when gene i has a fold change of 3 of expression in cell type j, with respect to the overall mean expression within that gene and cell type.

We assess the significance (Fisher’s exact test) of the overlap between all four ROSMAP network predictions and single cell observations, for each cell type separately. In each comparison, we remove known gene markers to avoid optimistic and unfair overlap assessments. To enable the comparison between ROSMAP and GMSCA cell types, we aggregated excitatory and inhibitory neurons into a single type. For the rest, we used the exact equivalent between the two sources (e.g. we did not consider oligodendrocyte precursors as oligodendrocytes).

### Using EWCE to replicate cell-specific expression enrichment from GSMCA predictions

EWCE was used to assess whether each set of genes with multifunctional predictions for a specific cell type and network made by GMCNA show any enrichment in expression in an equivalent cell type from mouse. We should expect that when a set of genes are predicted to play any functional role at cell type ct, they should show reasonable values of expression in that cell type, even in different mammals (i.e. mus musculus in this case). We tested each predicted cell type gene set from GMCNA in each network against enrichment in the equivalent cell type at EWCE. For such purpose we used the bootstrap.enrichment.test() R function from the EWCE package, but firstly we converted the human genes coming from our networks into MGI symbols using the mouse_to_human_homologs data table from EWCE. It provides the MGI mouse gene symbol for each HGCN human gene identifier. Gene markers were removed from the analysis. We used all MGI genes at the mouse_to_human_homologs table as the background gene set and 10000 permutations for each test. We were just interested in level 1 annotations. Results with P < 0.05 were reported as significant.

## Resources

The software developed within this paper can be installed from

https://github.com/drlaguna/GMSCA

The software used to develop primary co-expression networks can be installed from

https://github.com/juanbot/CoExpNets

The PGCNs used in this paper are browasable at

https://snca.atica.um.es/coexp/Run/Catalog/

The three network families can be installed in the following three GitHub repositories

https://github.com/juanbot/CoExpGTEx

https://github.com/juanbot/CoExpROSMAP

https://github.com/juanbot/CoExp10UKBEC

## List of Figures & captions

**Supplementary Figure 1.**
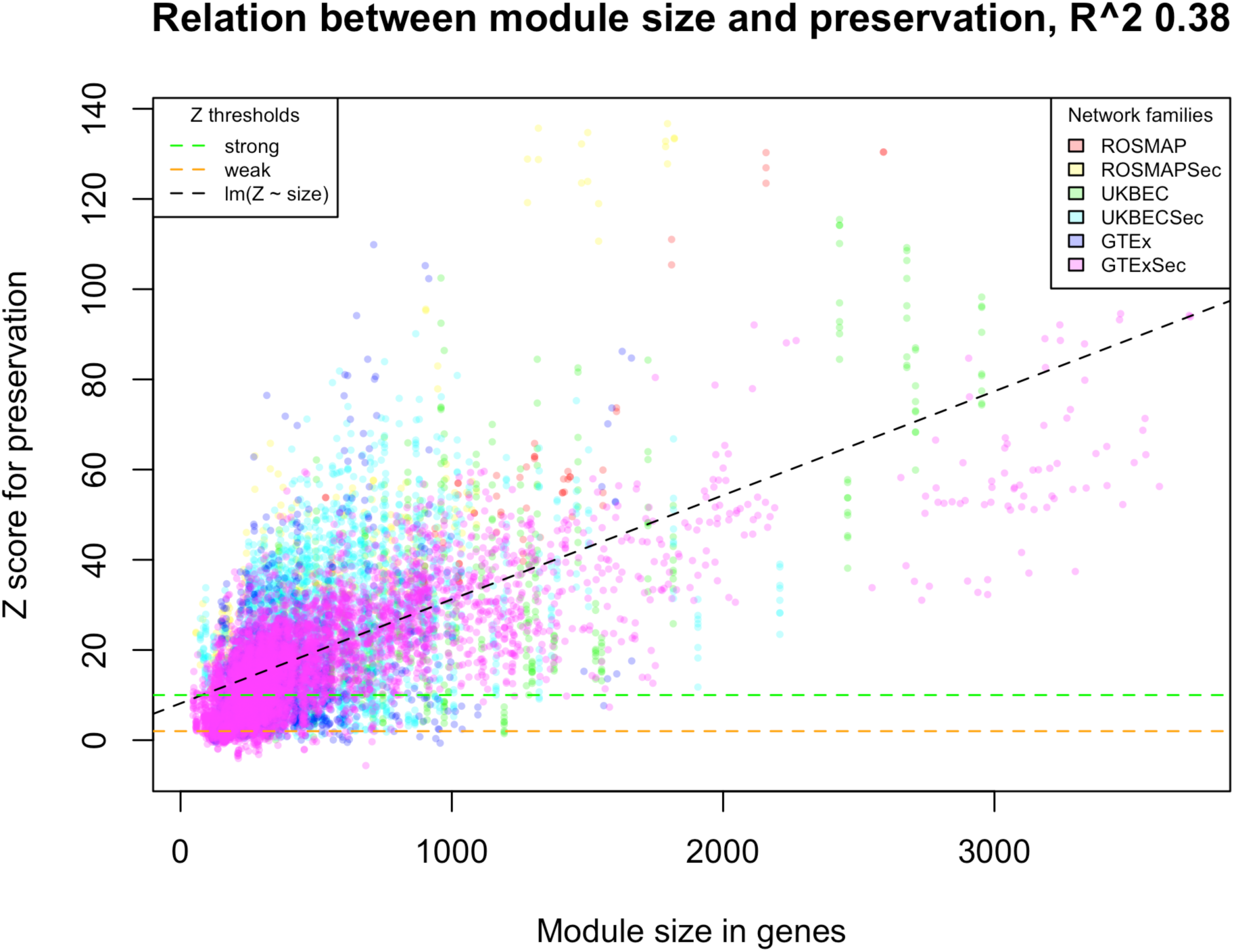
we observe a clear association between module size and preservation Z-score with an R2 of 0.38 in average for all tests we performed.

**Supplementary Figure 2.**
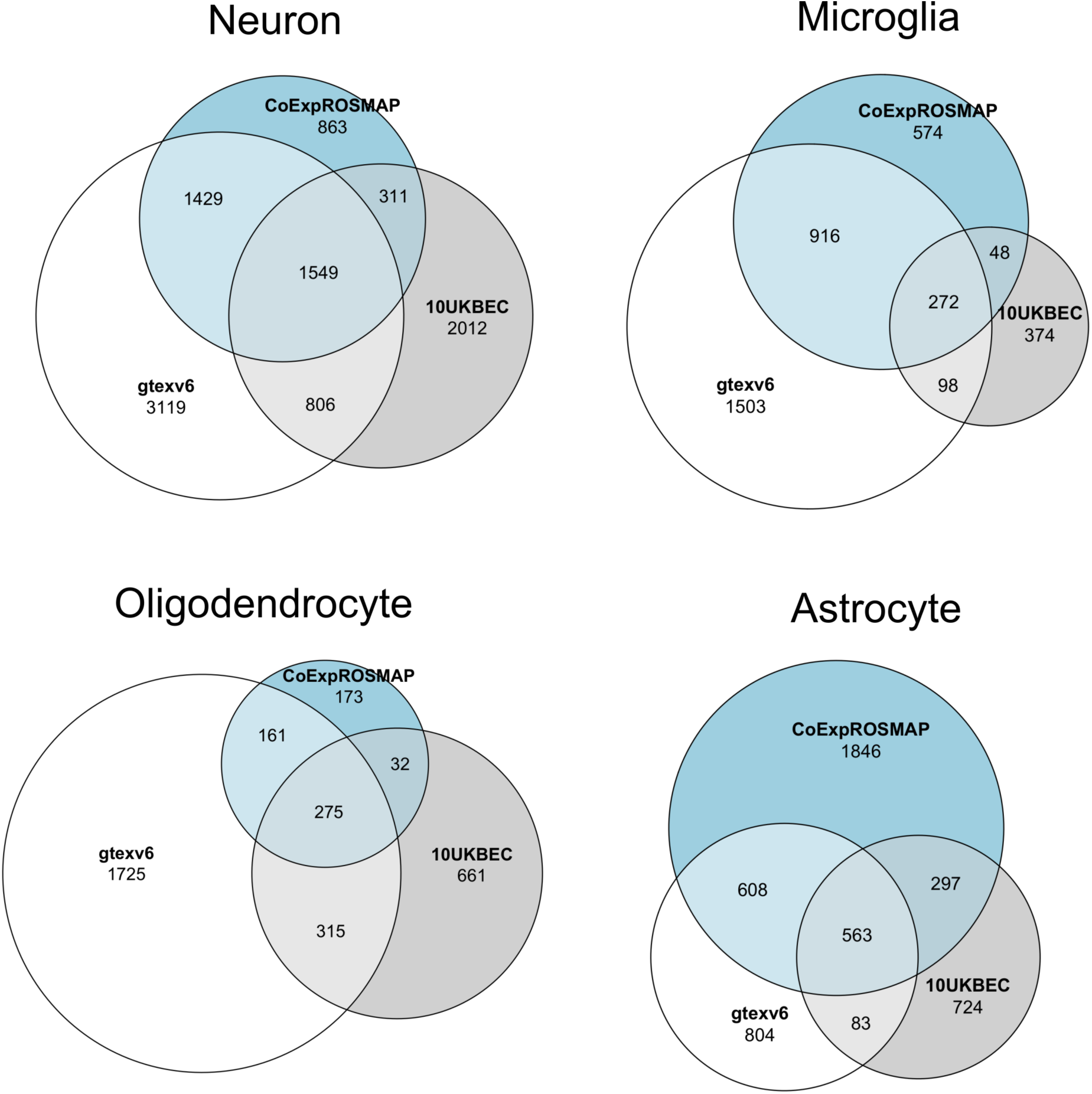
GMSCA predictions overlap between the three frontal cortex networks across cell types.

**Supplementary Figure 3.**
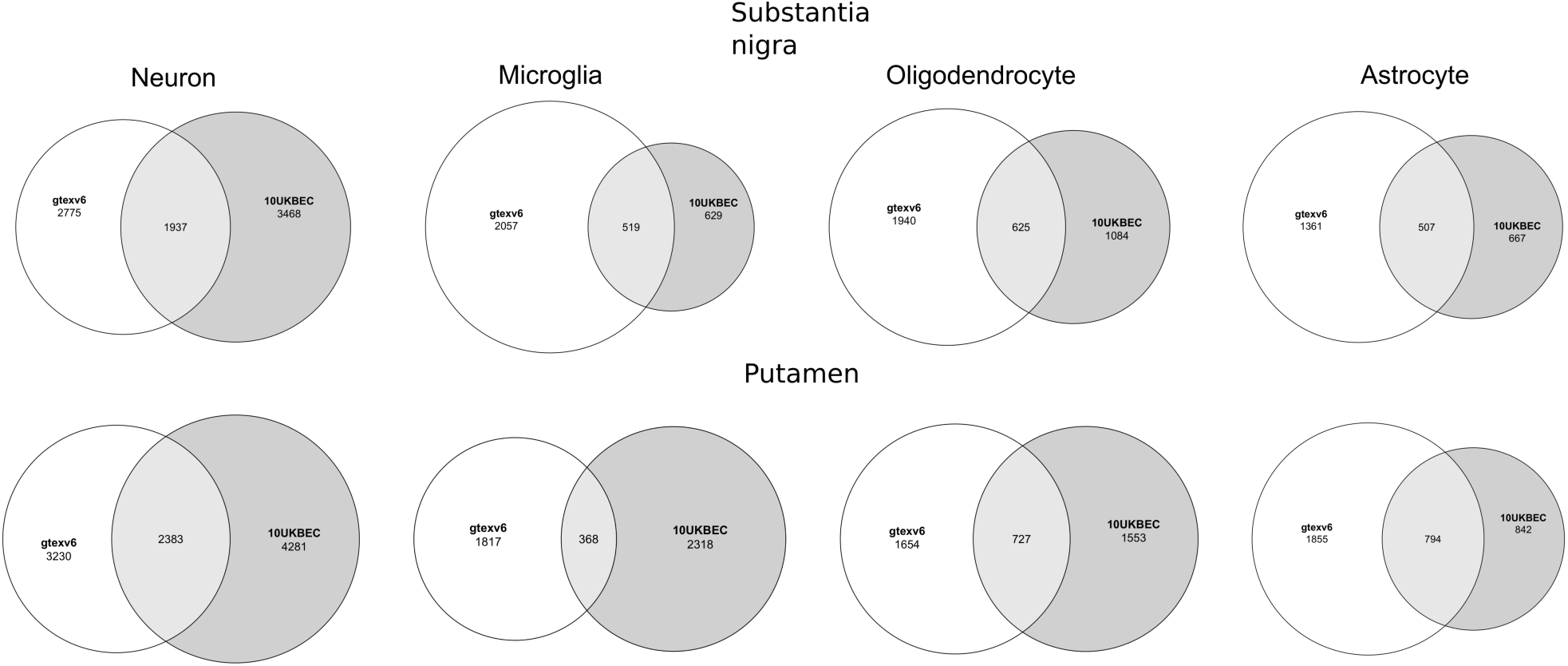
GMSCA predictions overlap between substantia nigra and putamen networks between UKBEC and GTEx

**Supplementary Figure 4.**
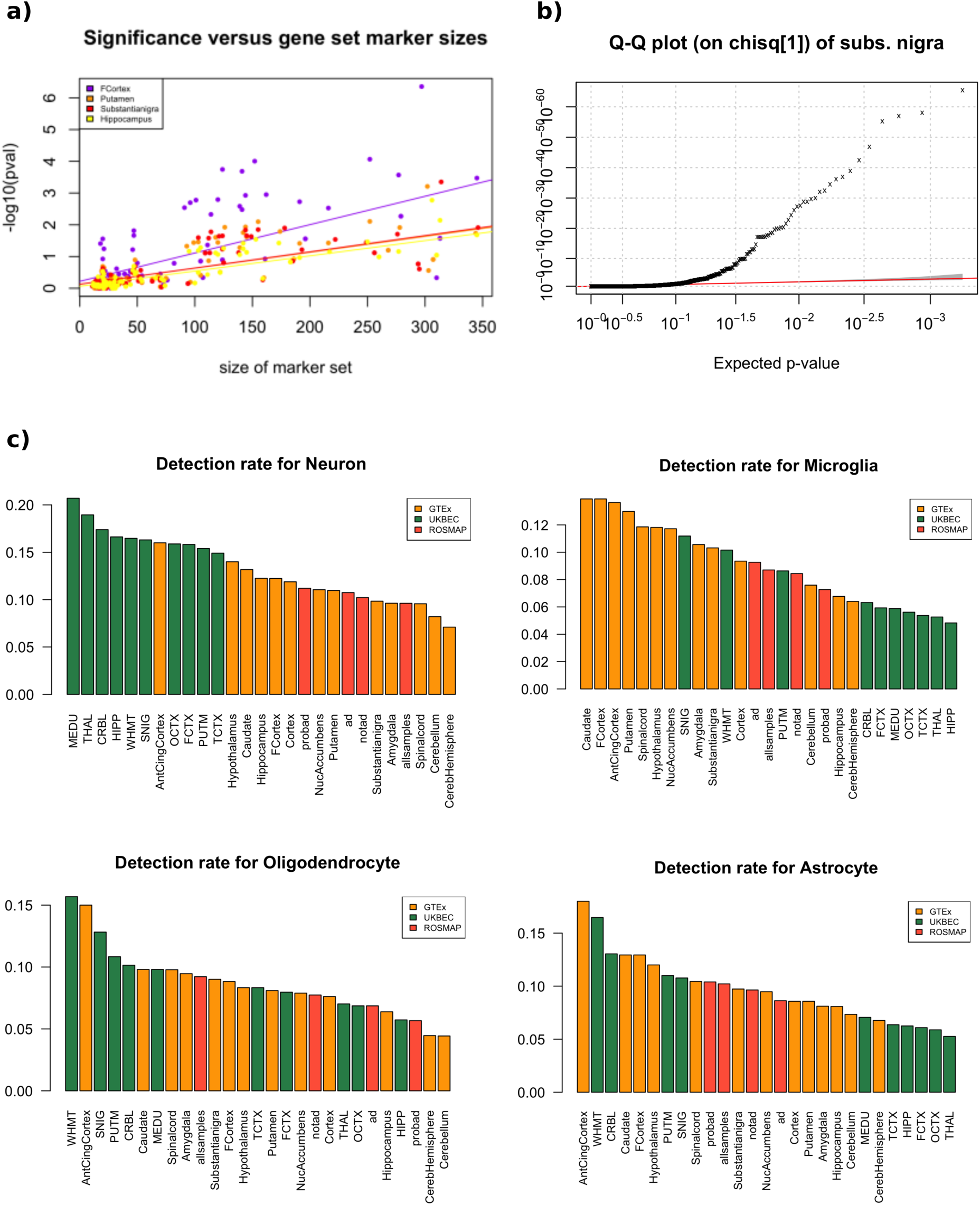
a) There is an association between the size of the gene marker sets employed at the cell enrichment tests (x-axis) and the p-value significance (y-axis), highest p-value 3.24E-6 for hypothalamus, mean R^2^ 0.28, min 0.17, max 0.45. We illustrate the association including just 4 tissues out of the 27 primary co-expression networks used in the paper. b) A Q-Q plot on the expected (Chi2 with 1 df) and observed p-values which clearly shows an inflation, substantia nigra is here used as an example. c) After we apply multiple testing Bonferroni correction within the network and regress out the module size effect from the resulting p-values, the new Q-Q plot shows no inflation. d) Decrease in enriched modules detected before correcting (red) and after Bonferroni plus module size regression (blue) for all 27 tissues in the paper.

## Tables

**Supplementary table 1.** Genes related to Parkinson Diseases in substantia nigra module from primary co-expression network (OMIM and Orphanet).

| **Disease** | **Gene** |
| --- | --- |
| PARKINSON DISEASE, LATE-ONSET; PD | NR4A2 |
| PARKINSON DISEASE, LATE-ONSET; PD | ATXN8OS |
| PARKINSON DISEASE 1, AUTOSOMAL DOMINANT | SNCA |
| PARKINSONISM-DYSTONIA, INFANTILE, 1; PKDYS1 | SLC6A3 |
| PARKINSON DISEASE 20, EARLY-ONSET; PARK20 | SYNJ1 |
| PARKINSONISM-DYSTONIA, INFANTILE, 2; PKDYS2 | SLC18A2 |
| Young-onset Parkinson disease | UCHL1 |

## Funding

R.H.R. was supported through the award of a Leonard Wolfson Doctoral Training Fellowship in Neurodegeneration. J.H. and M.R. were supported by the UK Medical Research Council (MRC), with J.H. supported by a grant (MR/N026004/) and M.R and S.G.R. through the award of a Tenure Track Clinician Scientist Fellowship (MR/N008324/1). J.H. was also supported by the UK Dementia Research Institute, The Wellcome Trust (202903/Z/16/Z), the Dolby Family Fund, and the NIHR. A.C. is supported by Fundación Séneca - Science and Technology Agency of the Region of Murcia, (grant reference: 20762/FPI/18). A.L.G.M is funded by Fundación Séneca (grant reference: 21230/PD/19).

## Acknowledgements

Study data to create the ROSMAP networks were provided by the Rush Alzheimer’s Disease Center, Rush University Medical Center, Chicago. Data collection was supported through funding by NIA grants P30AG10161, R01AG15819, R01AG17917, R01AG30146, R01AG36836, U01AG32984, U01AG46152, the Illinois Department of Public Health and the Translational Genomics Research Institute.

## Notes

### Competing Interest Statement

The authors have declared no competing interest.

